# Reward processing electrophysiology in schizophrenia: Effects of age and illness phase

**DOI:** 10.1101/2020.06.18.158469

**Authors:** Samantha V. Abram, Brian J. Roach, CB Holroyd, MP Paulus, Judith M. Ford, Daniel H. Mathalon, Susanna L. Fryer

## Abstract

**Background:** Reward processing abnormalities may underlie characteristic pleasure and motivational impairments in schizophrenia. Some neural measures of reward processing show strong age-related modulation, highlighting the importance of considering age effects on reward sensitivity. We compared event-related potentials (ERPs) reflecting reward anticipation (stimulus-preceding negativity, SPN) and evaluation (reward positivity, RewP; late-positive potential, LPP) across individuals with schizophrenia (SZ) and healthy controls (HC), with an emphasis on examining effects of chronological age, brain age (i.e., predicted age based on neurobiological measures), and illness phase.

**Methods:** Subjects underwent EEG while completing a slot-machine task for which rewards were not dependent on performance accuracy, speed, or other preparatory demands. Slot-machine task EEG responses were compared between 54 SZ and 54 HC individuals, ages 19 to 65. Reward-related ERPs were analyzed with respect to chronological age, categorically-defined illness phase (early; ESZ versus chronic schizophrenia; CSZ), and were used to model brain age relative to chronological age.

**Results:** Illness phase-focused analyses indicated there were no group differences in average SPN or RewP amplitudes. However, a group x reward outcome interaction revealed that ESZ differed from HC in later outcome processing, reflected by greater LPP responses following loss versus reward (a reversal of the HC pattern). While brain age estimates did not differ among groups, depressive symptoms in SZ were associated with older brain age estimates while controlling for negative symptoms.

**Conclusions:** ESZ and CSZ did not differ from HC in reward anticipation or early outcome processing during a cognitively undemanding reward task, highlighting areas of preserved functioning. However, ESZ showed altered later reward outcome evaluation, pointing to selective reward deficits during the early illness phase of schizophrenia. Further, an association between ERP-derived brain age and depressive symptoms in SZ extends prior findings linking depression with reward-related ERP blunting. Taken together, both illness phase and age may impact reward processing in SZ, and brain aging may offer a promising, novel marker of reward dysfunction that warrants further study.

## Introduction

Reward processing deficits are a core feature of schizophrenia (SZ) (1–4) that may underlie characteristic motivational impairments (3) and contribute to poor functional outcomes (5–7). Many reward-focused studies i) implement paradigms that confound reward processing with other cognitive or behavioral demands known to be impaired in SZ (4,8), ii) do not focus on key demographic variables (like age) (9), or iii) do not consider clinical variables that may be associated with accelerated brain changes (like illness phase) (10).

Prior research demonstrates impairments in SZ when anticipating future reward enjoyment, but suggests that “in-the-moment” consummatory, pleasure may be relatively intact (based on experience sampling, behavioral performance, and functional MRI) (11,12). With millisecond resolution, event-related potentials (ERPs) can capture temporally distinct reward processing phases. We examined three sequential reward-related ERP components that span the anticipatory and consummatory reward phases: the stimulus preceding negativity (SPN, reflecting reward anticipation), the reward positivity (RewP, reflecting early reward evaluation), and the late positive potential (LPP, reflecting later, and therefore higher-order, aspects of reward evaluation).

The SPN builds several hundred milliseconds prior to an anticipated stimulus (13,14). In the context of a reward task, the SPN is sensitive to reward anticipation, i.e., larger (more negative) when expecting reward versus non-reward (15,16). Although the SPN has not been a focus of many reward studies in SZ, aberrant SPN responses to certain versus uncertain reward-signaling cues has been demonstrated in SZ (17).

The RewP is a positive medial frontal component occurring ∼250–300 ms after reward feedback relative to non-reward or loss (18). Historically, research focused on the negativity following loss, typically referred to as the feedback-related negativity (FRN) (19–21), often calculated as loss – win difference; although more recent work emphasizes win-related contributions to RewP variance (22–24) and promotes calculation of the difference score to reflect a positivity from rewards (i.e., by measuring a win – loss difference wave, rather than the originally measured loss – win) (18). Blunted RewP is thought to reflect reduced reward sensitivity and is evident in those with, or at-risk-for, major depressive disorder (MDD) (18). Studies using passive reward tasks reported intact RewP in SZ (25,26), though a study of probabilistic learning found attenuated RewP correlated with more severe negative symptoms (27), raising the possibility that early reward feedback processing (i.e., “consummatory”) deficits are most evident among SZ with prominent negative symptoms.

The LPP is a slow, centro-parietal, ERP component typically measured starting ∼600 ms following stimulus onset, reflecting sustained engagement with salient content (28,29). The LPP follows the RewP in time, capturing later-stage outcome processing in reward tasks (30), and may reflect more downstream processing of output from the earlier evaluation systems (31). The LPP has frequently been examined using emotion-picture viewing tasks (28,32–34), though more recent studies demonstrated the LPP’s role in reward processing in healthy controls (HC) (35–39) and motivated attention to potential rewards or punishments in SZ (40). Blunted LPP following reward gains has also been associated with higher negative symptoms in a transdiagnostic sample that included schizophrenia-spectrum disorders (41).

Importantly, normal and pathological aging effects may impact reward processing neural signals. RewP magnitude shows a negative association with age among HC (9), and a depression-focused meta-analysis found that RewP blunting was most pronounced in younger depressed samples (42). Beyond reward processing, growing evidence supports hypotheses of accelerated brain aging in SZ (43–47), particularly early in the illness course (44,46), which may be due to abnormal brain maturation (48). Brain aging has been measured as the difference between chronological and predicted age based on neurobiological measures (referred to as “BrainAGE” gap). Most of the BrainAGE literature to date has used structural MRI data to estimate age, though this framework is expanding to other neurobiological measures, including a recent study using resting EEG (49). Incorporating neural age estimates may enhance sensitivity for detecting age-associated pathophysiological processes in psychiatric disorders (50). Given age relationships for some reward ERPs and the importance of motivational and reward processes in normal neurodevelopment (51), examining brain aging relationships in SZ with ERP reward-processing metrics is warranted.

SZ and depression are highly comorbid disorders (52,53). The presence of reward processing deficits among individuals with SZ and depressive disorders has led researchers to characterize the neural processes linking common symptoms across these disorders, like anhedonia (2). While there is a sizeable literature on the RewP and depressive symptoms and risk for depression (18,54), no studies have examined RewP in SZ as a function of depressive features (55). Accordingly, there is strong empirical impetus to evaluate relationships between RewP and depressive symptoms in SZ.

Here we use a slot-machine task in which rewards were not dependent on decision-making, response speed, or performance accuracy (26), to isolate reward processes from other preparatory and/or executive demands (56). Each trial depends on the pseudorandom population of three reels with fruit symbols. The reels sequentially populate, left to right, enabling parsing of anticipatory from early and late-consummatory signals. Because neurodevelopmental changes are central to SZ pathogenesis and healthy reward processing (51,57), we compared SZ and HC across a wide age range, with the goal of examining contributions of i) age through analyses of chronological age and BrainAGE (derived from reward ERPs), and ii) categorically-defined illness phase through recruitment of early illness (ESZ) and chronic (CSZ) patient groups (9,10).

We expected blunting of anticipatory signals (SPN) based on current models of reward deficits in schizophrenia (11), and more pronounced group differences during later reward evaluation (LPP) given theories that downstream processes following initial feedback may be integral to reward dysfunction in SZ (4). Based on prior electrophysiology studies, we did not expect group differences in early reward evaluation indexed by the RewP (25,26), but did expect reduced reward-related ERP amplitudes (SPN, RewP, LPP) to correlate with greater negative symptoms (27,41). We also directly compared reward-anticipation with reward-outcome ERPs across the groups, as a prior report found that anticipatory signals (SPN) correlated with later outcome processes (LPP) in HC (37). Finally, we hypothesized depressive features would relate to RewP amplitude, based on growing evidence that blunted RewP is a marker of depression vulnerability (18), and because depressive symptoms frequently occur in SZ (52).

## Methods and Materials

### Subjects

Fifty-four SZ (76% men; age range = 19.07–64.70 years) and 54 HC (78% men; age range = 19.25–64.41 years) were recruited via community advertisements; results from the HC sample are described in a previous study (58). SZ were either early illness (ESZ) within 5 years of onset (59–61) (*n* = 26; mean illness duration = 2.90 ± 1.47 years), or chronic (CSZ; *n* = 28; mean illness duration = 23.55 ± 15.35 years). ESZ and CSZ were comparable in gender representation, handedness, chlorpromazine equivalents (CPZeq), haloperidol equivalents (HPeq; 56), and concomitant anti-depressant treatment (see Supplemental Materials); and, as expected based on subgroup assignment, ESZ and CSZ differed in age and illness duration (Table 1).

**Table 1.** Sample Demographics

|  | HC (*n* = 54) | ESZ (*n* = 26) | CSZ (*n* = 28) | F/t |
| --- | --- | --- | --- | --- |
| **Demographics** |  |  |  |  |
| Age, years | 33.72 (14.42) | 24.47 (4.01) | 44.05 (14.73) | 15.77*** |
| Gender (% male) | 77.78 | 73.08 | 78.57 | 0.14 |
| Handedness (% right) | 87.04 | 92.31 | 82.14 | 0.43 |
| Illness Duration, years | --- | 2.90 (1.47) | 23.55 (15.35) | 6.71*** |
| <sup>a</sup>CPZeq, mg | --- | 301.65 (257.85) | 386.47 (304.70) | 0.98 |
| <sup>b</sup>HPeq, mg | --- | 7.06 (6.77) | 6.21 (3.45) | 0.19 |
| Anti-depressants (% sample) | --- | 38.46 | 28.57 | 0.58 |
| **Clinical Symptoms** |  |  |  |  |
| CAINS Motivation & Pleasure | --- | 16.64 (7.44) | 14.48 (8.00) | -1.02 |
| CAINS Expression | --- | 4.35 (3.63) | 3.63 (4.43) | -0.64 |
| PANSS Total Negative | --- | 15.35 (4.45) | 15.73 (6.75) | 0.24 |
| PANSS Total Positive | --- | 16.27 (6.39) | 16.67 (4.97) | 0.25 |
| PANSS Depression | --- | 8.77 (3.67) | 7.79 (3.60) | 0.33 |

\* $p < .05$ ; \*\*\* $p < .001$
<sup>a</sup>Three subjects were taking first-generation antipsychotics, and 34 subjects were taking second-generation antipsychotics.
<sup>a</sup>CPZeq and HPeq were correlated at .85.
Abbreviations: *HC*, healthy controls; *ESZ*, early illness schizophrenia subjects; *CSZ*, chronic illness schizophrenia subjects; *CPZeq*, chlorpromazine equivalents; *HPeq*, haloperidol equivalents; *CAINS*, Clinical Assessment Interview for Negative Symptoms; *PANSS*, Positive and Negative Syndrome Scale.

We used the Structured Clinical Interview for DSM-IV (SCID-IV-TR) (63) to confirm a schizophrenia or schizoaffective diagnosis for SZ and to exclude HC if they met criteria for a past or current Axis I disorder. HC were also excluded for having a first-degree relative with a schizophrenia-spectrum disorder. Urine toxicology tested for common drugs of abuse (e.g., opiates, cocaine, amphetamines) and potential subjects with a positive test were excluded. English fluency was required for participation. Additional exclusion criteria for SZ and HC were history of head injury, neurological illness, or other major medical illness that impacts the central nervous system. Study procedures were approved by the Institutional Review Board at the University of California, San Francisco. All subjects provided written informed consent.

### Task description

We developed a 288-trial slot-machine reward task adapted from prior studies (64–66), and shown to elicit expected SPN, RewP, and LPP condition effects in the same HC sample studied here (58). Subjects initiated each trial via a button press, after which reels 1–3 (R1, R2, R3) populated automatically from left to right with single, sequential fruit symbols. After R3 populated, feedback indicated a win or loss. The *reward anticipation* phase spanned the population of R1 and R2 (culminating just prior to R3; 0–3666 ms); the *reward evaluation phase* began with the population of the R3 symbol (3666–6115 ms).

Trial types were wins, near misses, and total misses. Wins occurred when all three reels populated identical fruit symbols (AAA). Near misses occurred when R1 and R2 populated matching symbols but the R3 symbol was incongruent (AAB). Wins and near misses were congruent on R1 and R2, inducing similar reward anticipation just prior to R3. Total misses occurred when R2 did not match R1 (ABC), indicating a loss at R2 and eliminating further reward anticipation prior to R3 (with R3 providing no additional information about the trial’s outcome). To reflect real-world slot-machine outcomes, subjects encountered more frequent total misses (*n* = 144, P = .50) than wins (*n* = 72, P = .25) and near misses (*n* = 72, P = .25).

Wins yielded a $1.25 payout, while near and total misses yielded $0 payouts. Subjects were instructed that they would receive monetary compensation reflective of their slot-machine winnings, in addition to routine compensation for participation. Supplemental Materials contain additional task details.

### Clinical symptom ratings

Negative symptoms were evaluated using the Clinical Interview for Negative Symptoms (CAINS) (67) and the Positive and Negative Syndrome Scale (PANSS) (68). The CAINS is a newer negative symptom measure, with strong psychometric properties (69), that was designed to parse deficits in *motivation and pleasure* for social, vocational, and recreational activities (MAP; 9 items) from deficits in verbal and non-verbal *expression* (EXP; 4 items). From the PANSS, we computed a Total Negative symptom score (items 8-14) from the PANSS. We also used the PANSS to compute a Depression composite score based on a five-factor model that has been validated previously (70), and shown to correlate with several other depression measures in SZ (71,72); we summed the anxiety (G2), guilt (G3), and depression (G6) items to create the depression composite score (73).

### EEG acquisition and preprocessing

EEG data were recorded from 64-channel electrode cap using the BioSemi ActiveTwo system (www.biosemi.com). Data were digitized at 1024 Hz and a 0.1 Hz high-pass filter was applied using ERPlab (74). Reference electrodes were placed on the mastoids. Electrodes were placed above and below the right eye, and at the outer canthus of each eye, to record vertical and horizontal electrooculograms (VEOG, HEOG, respectively).

Data were entered into a modified version of the Fully Automated Statistical Threshold for EEG artifact Rejection (FASTER) pre-processing pipeline (75). This entailed: i) identifying outlier channels and interpolating their values in the continuous data, ii) removing outlier epochs from each subject’s trial set iii) applying spatial independent components analysis (ICA) to remaining trials, iv) identifying outlier components from spatial ICA using the ADJUST procedure (76), and v) removing outlier channels (see Supplemental Materials).

### ERP measurement

Epochs were time-locked to R3 onset and baseline corrected using the −100 to 0 ms preceding R3 for RewP and LPP, and −100 to 0 ms preceding R2 (−1300 to −1200 ms preceding R3) for SPN. A trimmed means approach excluded the top and bottom 10% at each time point before averaging to obtain a more robust estimate (77).

#### SPN (reward anticipation)

measured as the average voltage from −100 to 0 ms prior to the R3 outcome (14) from representative electrode Cz (66,78). We computed separate ERP averages for possible win trials (AA; collapsed across trials eventually revealed as wins (AAA) or near misses (AAB), as these are equivalent at R2) and total miss trials (AB; trials revealed as a loss at R2). For the SPN analyses, we removed two SZ subjects with average SPN values on total miss (AB) trials > 3 SD above the mean.

#### RewP (early reward evaluation)

measured as the average voltage from 228 to 344 ms post R3 onset; this time-window was chosen based on an average of measurements from a meta-analysis of 54 RewP studies (79). Statistical analyses were conducted based on representative electrode FCz (22,66,78,80). The RewP was computed as a difference score of wins minus near misses (AAA–AAB) (81); this isolates a pure valence effect as the stimulus probabilities are equated across these two conditions. We note that because the RewP is derived from a difference score, we cannot disentangle the contributions of wins and near misses.

#### LPP (late reward evaluation)

measured as the average voltage from 600 to 800 ms after R3 (82) from representative electrode Pz (30). We computed separate ERP averages for wins (AAA) and near misses (AAB). We removed four SZ subjects, from LPP analyses, with average LPP values on win (AAA) or near miss (AAB) trials > 3 SD above or below the mean.

### Data analysis

#### Age-adjusted ERP z-scores

We calculated age-adjusted z-scores for each ERP component to account for expected age differences between the clinical groups, ongoing neuromaturation processes expected in younger subjects, and normal aging processes expected in older subjects. This procedure removes normal aging effects while preserving variance associated with pathological age effects in SZ, similar to our prior ERP (83–85) and MRI studies (86,87).

To calculate age-adjusted z-scores, we first produced a HC age-adjusted regression model that regressed each ERP component onto age. We then extracted the intercept (*B*_0_), slope (*B*_1_), and root mean squared error (RMSE) from each model to compute the age-adjusted z-scores for all subjects, for the respective ERP component. We calculated age-adjusted z-scores for all subjects using the HC age-regression model:

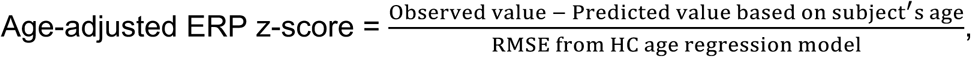

where the predicted value was calculated for each subject as follows:

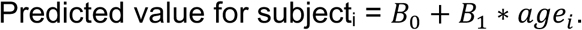

Accordingly, a given subject’s score reflects the deviation of their ERP component amplitude, in standard deviations units, from that expected for a HC of the same age.

#### Between-group ERP effects

We compared age-adjusted ERP measures across the three groups. For SPN, we used a mixed-effect ANOVA that included SPN z-scores as the outcome variable, Condition (AA, AB), Group (HC, ESZ, CSZ), and Group x Condition as fixed effects, and Subject as a random effect. We built an equivalent mixed-effects model for the LPP, but with Condition as AAA – ABC and AAB – ABC difference scores. For the RewP, we used a one-way ANOVA with RewP difference z-scores as the outcome variable and Group as a fixed effect. Follow-up tests were adjusted using Benjamini and Hochberg’s false discovery rate (FDR) algorithm (88).

#### Relationships between anticipatory and consummatory ERP components

Using three regression models, we tested if reward-anticipation was related to reward-outcome responses between HC and SZ. Each model included reward outcome ERPs (RewP, LPP AAA – ABC, LPP AAB – ABC) as the outcome variable, and reward anticipation (SPN AA – AB), Group (HC, SZ), and a Group X reward anticipation interaction term as the predictor variables.

#### ERP and negative symptom correlations

We correlated each ERP component of interest (SPN possible wins AA – AB, RewP wins AAA – near misses AAB difference scores, LPP wins AAA – ABC, and LPP near misses AAB – ABC) with negative symptoms (CAINS MAP, CAINS EXP, and PANSS Total Negative), yielding 12 correlations total; we adjusted for multiple comparisons using the FDR algorithm. Correlations were computed across all SZ as we were interested in dimensional associations regardless of illness phase.

#### RewP and depressive symptom correlation

We correlated RewP difference scores with the PANSS Depression composite score.

#### Brain age analyses

When deriving a reward-related BrainAGE model, we included the specific contributions of reward anticipation and outcome signals, while also accounting for more general motivational salience following an outcome (collapsing across wins and near misses). To isolate reward anticipation, we included the difference between SPN possible wins and total misses (AA – AB), and included consummatory measures of reward outcome processing with the RewP difference wave (wins AAA – near misses AAB) and the difference between LPP wins and near misses (AAA – AAB). We measured general motivational salience after an outcome as the average of LPP wins and near misses (AAA + AAB)/2. We then constructed the final BrainAGE model from unadjusted HC ERP data, including the four conditions described above. Results from this model are described in Supplemental Materials.

We used the HC regression weights from the BrainAGE model to predict BrainAGEs for all subjects, and then calculated the BrainAGE gap as BrainAGE minus chronological age (44). (One subject with a predicted BrainAGE > 3SD above the mean was removed from subsequent analyses.)

Lastly, we tested whether negative and depressive symptoms were related to the BrainAGE gap, based on evidence that accelerated aging is related to SZ symptoms (47); we regressed BrainAGE gap onto CAINS MAP, CAINS EXP, and PANSS Depression scores for all SZ (we did not include PANSS Total Negative given high collinearity with the CAINS; 60% of PANSS Total Negative variance was explained by the CAINS).

#### Effects of antipsychotic medication dosage and anti-depressant treatment

We evaluated pairwise correlations between CPZeq and HPeq with all ERPs, symptom, and BrainAGE measures. We also tested whether ERPs, symptoms, or BrainAGE metrics were related to concomitant anti-depressant treatment via *t*-tests.

## Results

### Reward-related ERP effects

Grand average ERP waveforms are presented in Figure 1 and RewP difference waveforms in Figure 2 (Supplemental Materials). As reported in (58), HC showed the expected condition effects for SPN, i.e., more negative for anticipated wins than total misses (AA < AB), as well as RewP and LPP, i.e., more positive for wins than near misses (AAA > AAB).

**Figure 1.**
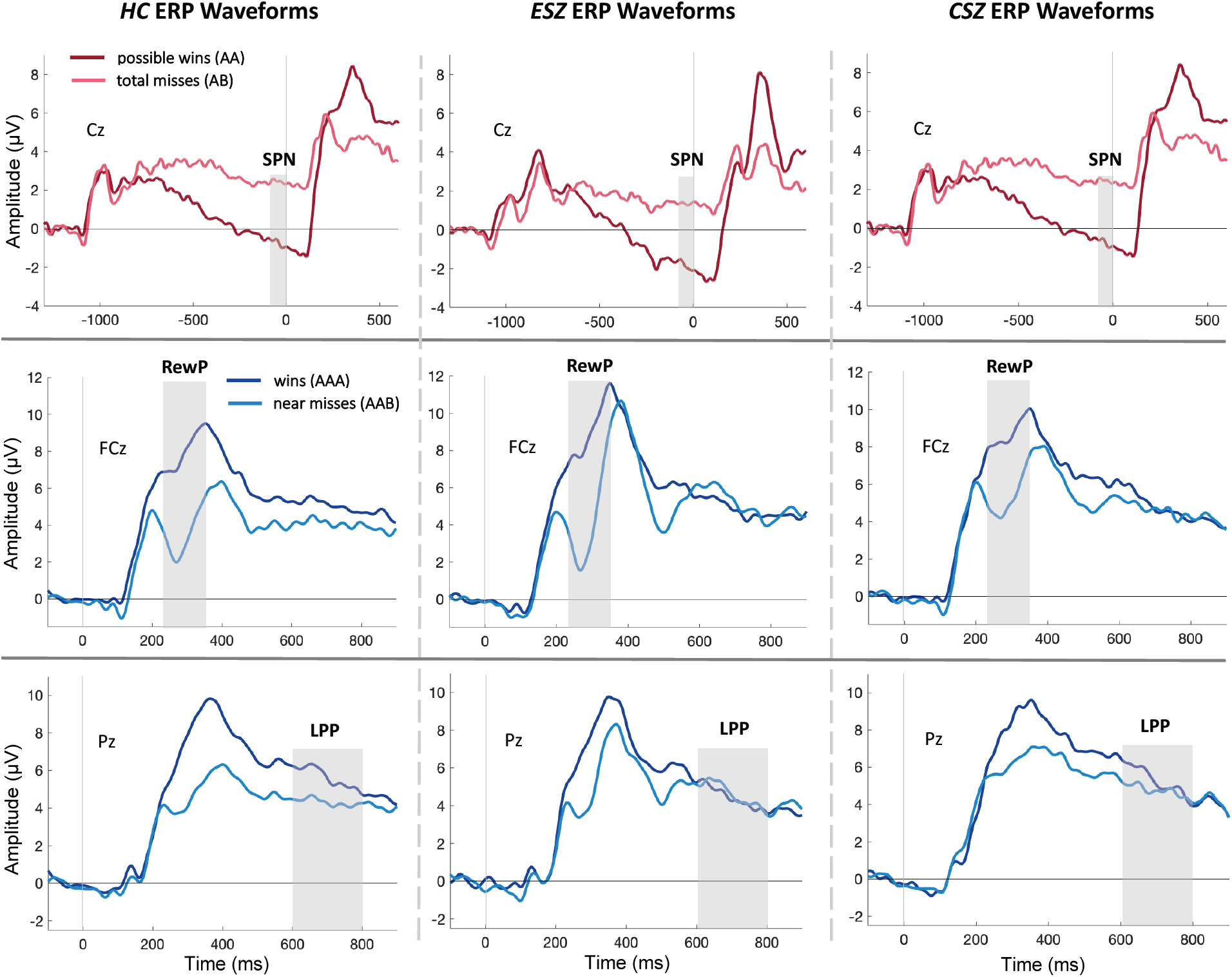
Grand average waveforms. Top: Stimulus preceding negativity (SPN) grand average waveforms at electrode Cz; win (AA) trials shown in red and total miss (AB) trials in pink. Middle: Reward Positivity (RewP) grand average waveforms at electrode FCz; win (AAA) trials shown in dark blue and near miss (AAB) trials in light blue. Bottom: LPP grand average waveforms at electrode Pz; win (AAA) trials shown in dark blue and near miss (AAB) trials in light blue. Time at −1200 ms corresponds to Reel-2 outcome (top row); time at 0 ms corresponds with Reel-3 outcome. Grey bars represent the ERP measurement window. Abbreviations: *HC*, healthy controls; *ESZ*, early illness schizophrenia subjects; *CSZ*, chronic illness schizophrenia subjects.

**Figure 2.**
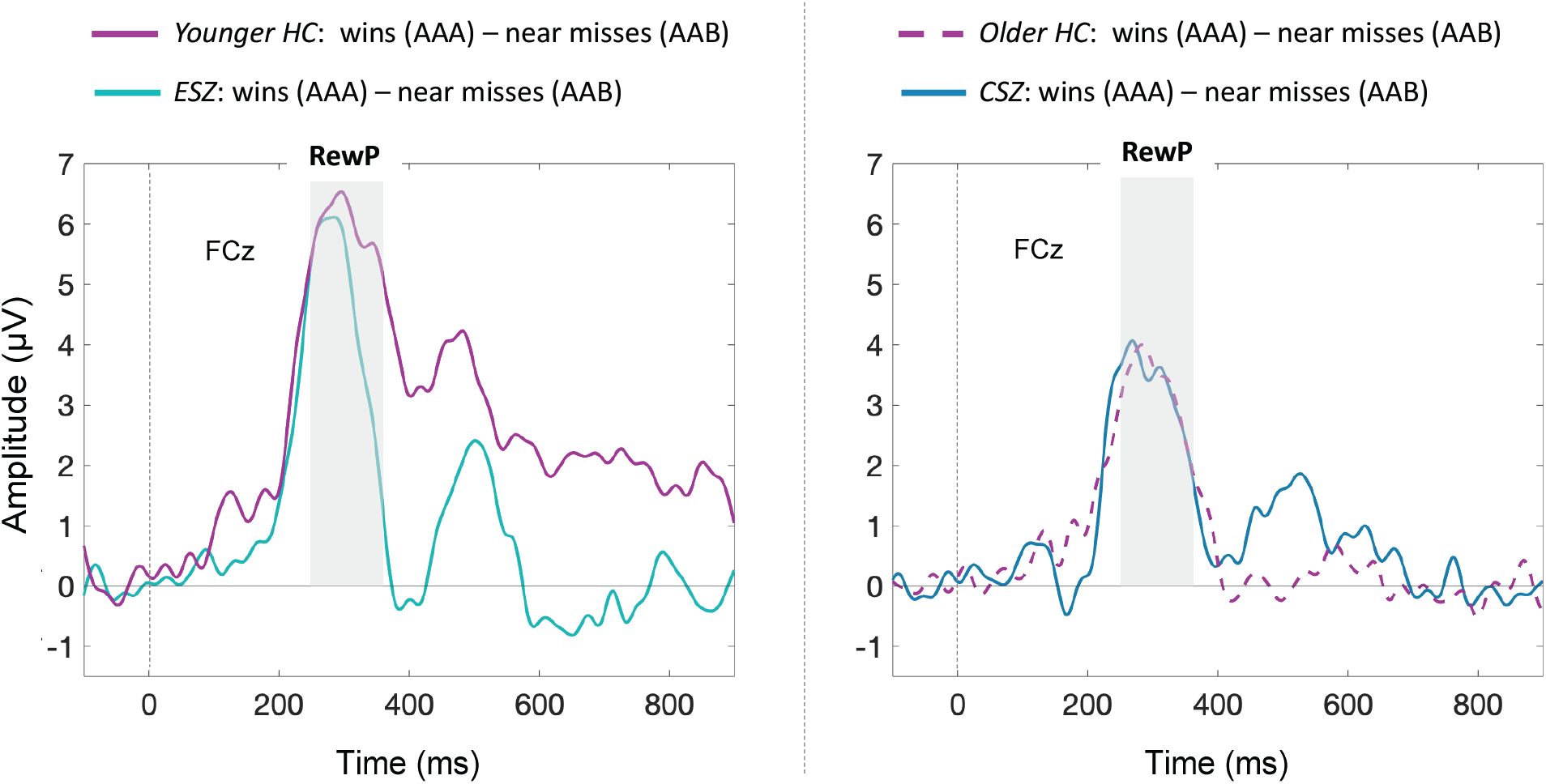
RewP difference waveforms. Difference ERP waveforms (wins AAA – near misses AAB) at electrode FCz. Time at 0 ms corresponds to Reel-3 outcome. Grey bar depicts reward positivity (RewP) measurement window. Healthy controls were divided using a median split, given a significant decline in RewP amplitude with age (Supplemental Materials). Abbreviations: *HC*, healthy controls; *ESZ*, early illness schizophrenia subjects; *CSZ*, chronic illness schizophrenia subjects.

#### SPN (reward anticipation)

There was no Group x Condition interaction for SPN z-scores (F_2,103_ = 0.55, *p* = .58), nor a main effect of Group (F_2,103_ = 2.07, *p* = .13; Figure 3B). All groups showed the expected AA < AB condition effect in the unadjusted data (Figure 3A).

**Figure 3.**
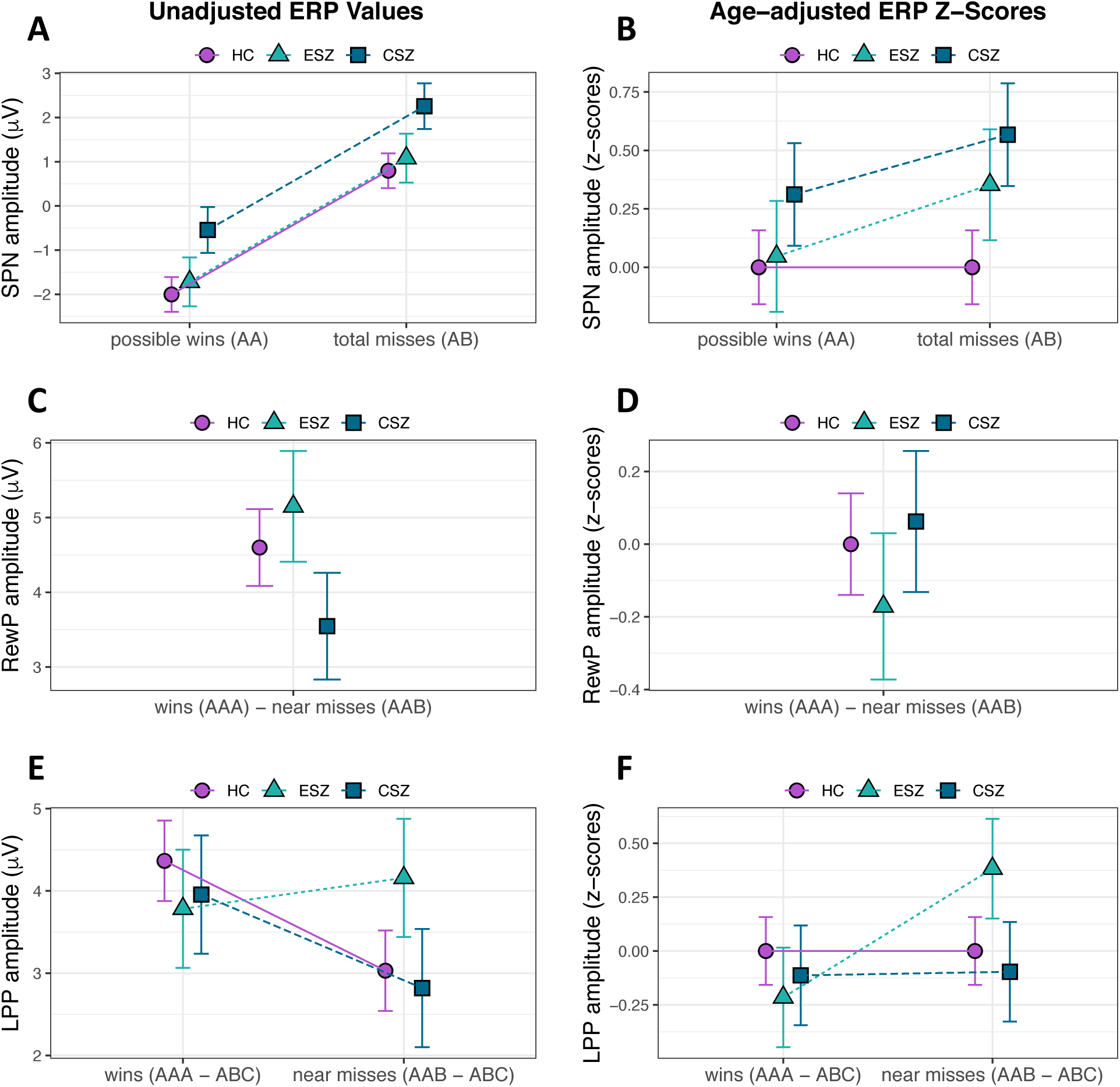
Group ERP effects. Results depicting group differences for the three ERPs; group means +/− standard error. (**A**) Stimulus preceding negativity (SPN) Condition effect for possible wins versus total misses regardless of Group (*HC*, healthy controls; *ESZ*, early illness schizophrenia subjects; *CSZ*, chronic illness schizophrenia subjects). (**B**) No Group x Condition interaction for age-adjusted SPN z-scores. (**C, D**) No Group differences for Reward Positivity (RewP) unadjusted or age-adjusted z-scores. (**E, F**) Significant Group x Condition interaction for late positive potential (LPP); whereby the ESZ group shows a heightened LPP response to near misses as compared to wins (**F**) when adjusting for HC age-related variance. For SPN analyses, we removed two SZ subjects with an average SPN value on AB trials > 3 SD above the mean; for the LPP analyses, we removed four SZ subjects with average LPP win (AAA – ABC) or near miss (AAB – ABC) values > 3 SD above or below the mean. Note for age-adjusted data: data were adjusted to account for normal aging effects using a z-scoring procedure based on a HC age regression model; as a result, HC z-scores have mean = 0 and SD = 1, and the patients group means reflect the degree and direction of abnormality, in standard units, from the HC-derived norms.

#### RewP (early reward evaluation)

There was no main effect of Group for RewP difference z-scores (F_2,105_ = 0.38, *p* = .68; Figures 3D, age-adjusted and Figure 3C, unadjusted).

#### LPP (late reward evaluation)

We observed a significant Group x Condition interaction for LPP z-scores (F_2,101_ = 3.76, *p* = .03): ESZ exhibited abnormally large LPP following near misses and abnormally small LPP following wins (AAA – ABC < AAB – ABC; *t*_101_ = – 3.18, *p* = .002, *p*_adj_ = .02; Figure 3F; Table S1), compared to the HC pattern of greater LPP responses to wins versus near misses (Figure 3E) (58), and the CSZ group, which showed no age-adjusted LPP condition difference).

The pattern of group effects for SPN, RewP, and LPP did not differ when using a HC age-matched grouping strategy in place of age-adjusted z-scores (see Supplemental Materials for full set of analyses).

### Anticipatory and consummatory ERP relationships

There was a significant Group interaction for RewP difference scores (F_1,102_ = 6.56, *p* = .01; Figure 4A), whereby SZ (*r*_50_ = −.53, *p* < .001) showed a negative relationship between SPN and RewP that was not observed among HC (*r*_52_ = −.01, *p* = .92). There was also a trend-level Group interaction for LPP wins (F_1,99_ = 3.69, *p* = .06; Figure 4B); here, both HC and SZ had a negative correlation between SPN and LPP wins (HC: *r*_52_ = −.31, *p* = .02, SZ: *r*_47_ = −.59, *p* < .001). For LPP near misses, there was a significant common slope across the groups (β = −.53, *p* < .001; Figure 4C), but no interaction (*p* > .10).

**Figure 4.**
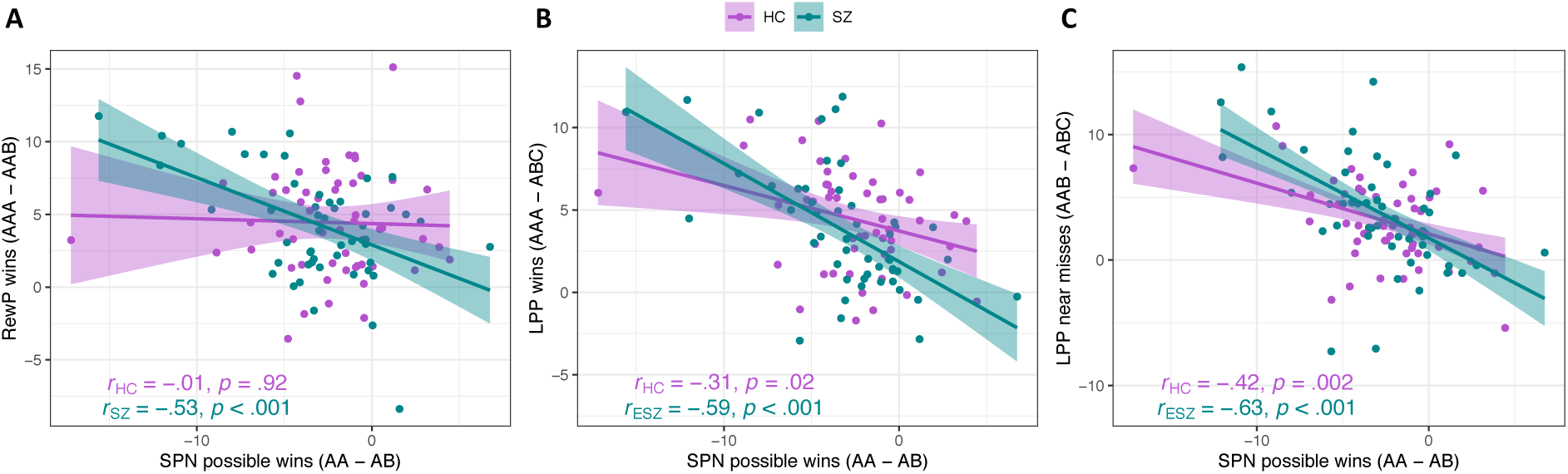
Relationships amonge reward ERP components. (**A**) Correspondence between SPN reward anticipation (AA – AB) with RewP difference (AAA – AAB) scores across groups (*HC*, healthy controls; *SZ*, schizophrenia subjects). (**B**) Correspondence between SPN reward anticipation with LPP win (AAA – ABC) scores across groups. (**C**) Correspondence between SPN reward anticipation with LPP near miss (AAB – ABC) scores across groups. For the SPN analyses, we removed two SZ subjects with an average SPN value on AB trials > 3 SD above the mean; for the LPP win analyses, we removed three SZ subjects with an average value > 3 SD above or below the mean; and for the LPP near miss analyses, we removed one SZ subject with an > 3 SD above the mean.

### ERP clinical symptom correlations

Across all SZ, negative symptoms were not correlated with any ERP measures (all *p* > .10). Depressive symptoms were unrelated to RewP difference scores (*r*_52_ = – .12, *p* = .38).

### BrainAGE predictions for HC and SZ

Table S1 shows the HC model used to produce BrainAGEs for all subjects, which indicated that Age was negatively associated with RewP, positively associated with LPP, and had a marginal relationship with SPN, controlling for all other measures. A *t*-test indicated that BrainAGE predictions, on average, were similar for HC and SZ (*t*_105_ = −.46, *p* = .65; Figure 5A), and both groups showed a positive relationship between chronological age and BrainAGE (HC: *r*_52_ = .58, *p* < .001; SZ: *r*_51_ = .30, *p* = .02).

**Figure 5.**
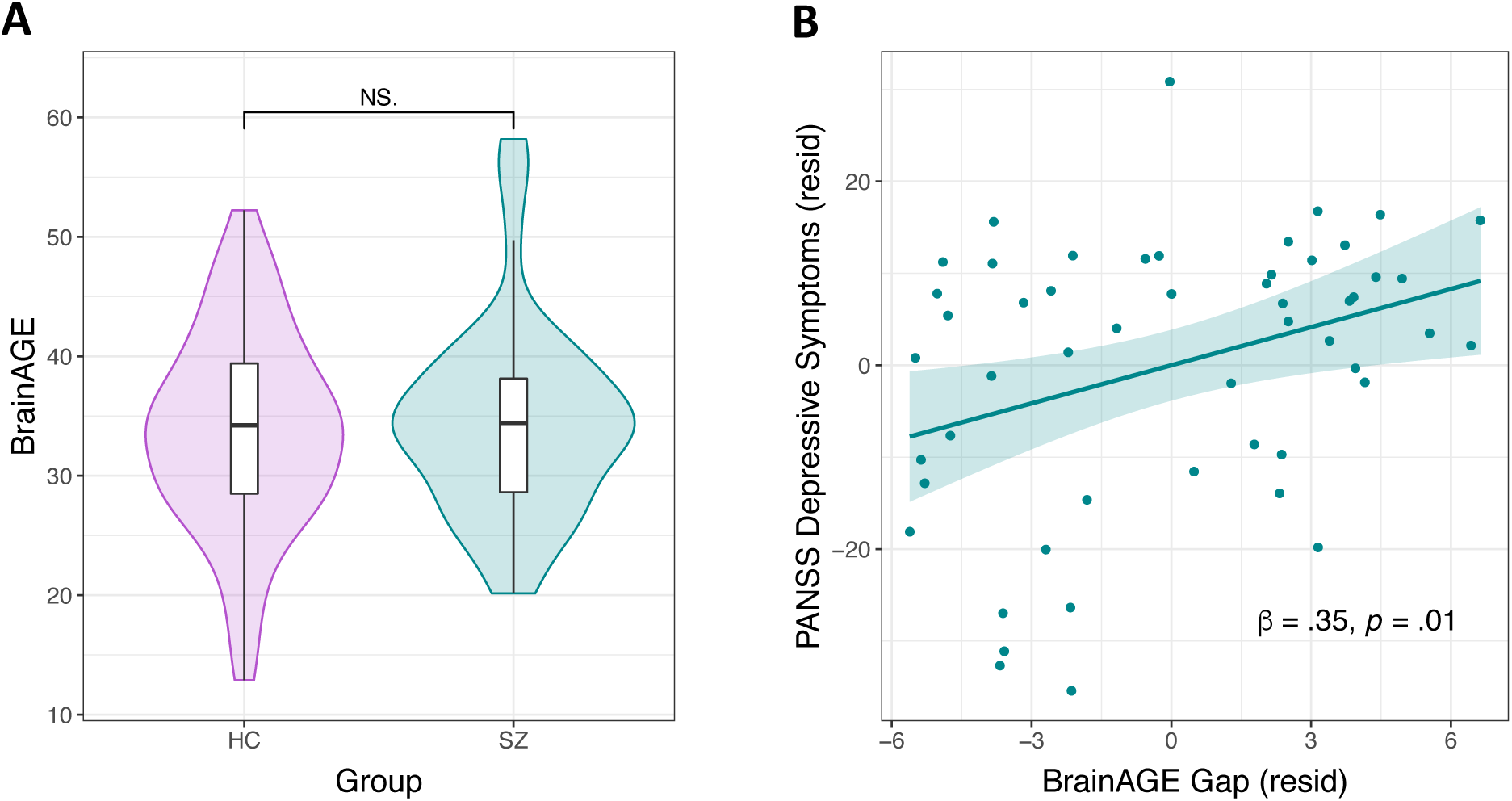
BrainAGE estimates and relationships with depressive symptoms. (**A**) Comparison of BrainAGE estimates (derived from reward ERPs) across groups (*HC*, healthy controls; *SZ*, schizophrenia subjects). (**B**) Relationship between depressive symptoms and BrainAGE gap scores in SZ; variables are residualized to account for the other predictors in Model 1 (Table 2). We removed one SZ subject with a BrainAGE estimate > 3 SDs above the mean from these analyses. Abbreviations: *PANSS*, Positive and Negative Syndrome Scale; *resid*, residuals.

### BrainAGE gap correlates with SZ depressive symptoms

Depressive symptoms were related to the BrainAGE gap among SZ when controlling for negative symptoms (β_depressive_symptoms_ = .35, *p* = .01; Figure 5B; Model 1, Table 2), while negative symptoms were unrelated to the BrainAGE gap; the bivariate correlation between the BrainAGE gap and depressive symptoms was also significant (*r*_51_ = .34, *p* = .01). Thus, the greater the estimated BrainAGE relative to an individual’s chronological age, the worse the depressive symptoms. Because depressive symptom ratings were higher for SZ with a schizoaffective diagnosis versus those with a schizophrenia diagnosis (*t*_52_ = 2.53, *p* = .01), and higher for SZ receiving concomitant anti-depressant treatment (*t*_52_ = 2.53, *p* = .01), we used hierarchical regression to test whether our BrainAGE-symptom effect was better explained by diagnosis type (schizophrenia or schizoaffective) or anti-depressant treatment (concomitant anti-depressant treatment versus not). Diagnosis and anti-depressant treatment did not predict the BrainAGE gap (Model 2, Table 2), account for the depressive symptom association, or improve the model (*R*^2^-change = 0.49, *p* = .61).

**Table 2.** Depressive symptoms correlate with BrainAGE gap in SZ

| | $\beta$ (SE) | t-stat | Adj-R<sup>2</sup> |
| --- | --- | --- | --- |
| **Model 1** |  |  |  |
| CAINS Motivation & Pleasure | .05 (.14) | 0.36 | .07 |
| CAINS Expression | .08 (.14) | 0.58 |  |
| PANSS Depression | .35 (.14) | 2.57* |  |
| **Model 2** |  |  |  |
| CAINS Motivation & Pleasure | .02 (.14) | 0.15 | .05 |
| CAINS Expression | .05 (.15) | 0.34 |  |
| PANSS Depression | .41 (.15) | 2.68* |  |
| Schizophrenia/Schizoaffective Diagnosis | -.09 (.33) | -0.29 |  |
| Anti-depressant Treatment | -.14 (.15) | -.93 |  |

\* $p < .05$
Model 1: $F_{3,48} = 2.35$ $p = .08$
Model 2: $F_{5,46} = 1.57$ , $p = .19$
Abbreviations: *CAINS*, Clinical Assessment Interview for Negative Symptoms; *PANSS*, Positive and Negative Syndrome Scale.

### Effects of antipsychotic medication dosage and anti-depressant treatment

CPZeq and HPeq were unrelated to any ERPs, symptom measures, or BrainAGE measures (all *p* > .10). Concomitant anti-depressant treatment was unrelated to any ERPs, negative symptoms, or BrainAGE measures (all *p* > .10).

## Discussion

We compared ERP components reflecting reward anticipation and evaluation in SZ versus HC, with a focus on investigating effects of chronological age, BrainAGE, and illness phase. ESZ exhibited aberrant late-stage reward evaluation signals relative to HC, indicated by heightened LPP responses following near misses versus wins. In comparison, reward anticipation and early reward outcome signals were intact in SZ. The relationship between reward anticipation and later outcome evaluation signals was also preserved in SZ (although SZ showed a relationship between reward anticipation and early reward processing that was not present in HC). BrainAGE predictions were derived from HC reward ERPs: while brain age estimates did not differ among HC and SZ, accelerated brain aging in SZ correlated with worse depressive symptoms, even after controlling for negative symptoms, schizoaffective diagnosis, and anti-depressant treatment. Together, our findings reveal novel age- and illness-phase indicators of altered SZ reward processing.

Contrary to our hypotheses, SZ did not differ from HC in their SPN amplitudes, and SPN amplitudes were unrelated to negative symptoms. This suggests that the SPN may be relatively preserved in SZ during basic incentive processing. It is possible that anticipatory reward differences may emerge for SZ under more complex conditions, such as evaluating reward cues of varying certainty (17). This follows from interpretations that certain reward deficits stem from interactions with higher-order cognitive processes like attention or working memory (26,89–91), and from evidence that avolition and anhedonia are linked to learning and other higher-order processes (92). Effects of negative symptoms on reward anticipation have also been observed in tasks where cues indicate the need to prepare a response to either win, or avoid losing, different amounts of money, like in the monetary incentive delay task (93,94). Taken together, the circuitry underlying passive reward anticipation (95) may be relatively intact in schizophrenia, whereas deficits in performance-based reward anticipation circuits may have implications for negative symptoms; although this needs to be directly tested by contrasting these two reward types in the same study.

RewP responses for reward versus non-reward outcomes were comparable for SZ and HC during a simple reward task, replicating earlier case-control studies that found intact RewP among SZ (25,26). This finding also fits with broader evidence for intact immediate consummatory responses in SZ (11). However, when comparing slopes of SPN reward anticipatory signals with early reward outcome processing measured by RewP, we found a relationship in SZ that was not observed in HC: that is, more reward anticipation (i.e., more negative SPN) was associated with a larger immediate response to reward (i.e., more positive RewP) in SZ. This could suggest that SZ have a unique temporal coupling between these reward signals that differs from HC; although it is unclear whether this is a disease-driven correlation or a typical relationship that was not present in our HCs. In comparison, both HC and SZ showed a significant SPN – LPP relationship, consistent with a prior report from a HC sample during a rewarded time estimation task (37).

Illness phase differences emerged during later reward outcome processing. More specifically, ESZ had larger LPP responses following near misses than wins, a reversal of the HC pattern (58). This alteration was specific to ESZ, as the CSZ group showed no age-adjusted LPP differences. One potential explanation for these findings could be differences in attention allocation during reward processing. The LPP is modulated by attention and can be reduced if attention is directed away from emotionally-arousing content (97). Greater LPP in ESZ following near miss events could reflect more focus on negative outcomes or insufficient focus to positive outcomes, or both. Alternatively, HC could be suppressing their LPP to near misses by paying less attention to unrewarding outcomes. HC might also be enhancing their LPP to wins by paying more attention to rewarding outcomes, or by enhancing their anticipation of the next trial. This fits with interpretations that the LPP indicates reward-related attentional deployment; for instance, one study found that LPP amplitudes were increased if subjects directed their attention towards cues indicating monetary earnings (98).

Finally, when collapsing across illness phase, accelerated brain aging correlated with higher depressive symptoms, when controlling for negative symptoms and schizoaffective diagnosis; i.e., a higher BrainAGE relative to one’s chronological age was associated with worse depressive symptoms in all SZ. There is a wealth of literature linking reward responsiveness, particularly RewP, with an MDD diagnosis, self-reported depressive symptoms, and anhedonia (54,99–101). Our finding adds to growing evidence relating reward responsiveness with age and depression liability (42), and highlights increasing efforts to elucidate dimensional markers of reward processing dysfunction that cut across traditional diagnoses, like MDD and SZ (2). This finding also underscores that brain-predicted aging measures can be used to capture meaningful variation in illness attributes within a disease group (102).

### Limitations

Though we found no relationships with chlorpromazine equivalents, medication status could still influence results. The ERP components we assessed were unrelated to negative symptoms, and we might have observed the hypothesized negative symptom relationships had we oversampled individuals meeting deficit syndrome criteria (103); it is also possible that these null results are due to our imperfect symptom measures (104), and/or failures to parse primary versus secondary negative symptoms (the latter which are caused by positive symptoms, treatment side effects, depression, or substance use). Future studies are needed to consider more inclusive BrainAGE models (e.g., adding multiple neurobiological modalities and extending the age range) that might improve prediction of normal and pathological reward processing trajectories, as we focused exclusively on ERP-derived reward-related metrics. We measured depressive symptoms using a scale developed for SZ and did not assess HCs, preventing dimensional analysis. A final caveat is that, while age and illness phase both modulated reward-related brain functioning, these features are inextricable in our cross-sectional samples, in that younger patients tend to be earlier in their illness course. Thus, longitudinal studies are needed to clarify the nature of the age and illness phase reward relationships suggested by our data.

### Conclusions

Our data highlight the impact of age, illness phase, and depressive features on reward-related brain functioning in SZ. Using a cognitively undemanding reward task, we identified areas of preserved functioning and illness-phase specific deficits: more specifically, reward anticipation (SPN) and early (RewP) reward evaluation were intact in ESZ and CSZ, while aberrant late-stage (LPP) reward evaluation emerged as a selective deficit among ESZ. SZ also showed more co-variation between reward anticipatory and outcome ERPs. Lastly, accelerated brain aging correlated with higher depressive symptoms across all SZ, extending prior findings linking depressive features and blunted RewP to the schizophrenia spectrum.

## Acknowledgments

Research supported by VA CX001028 to S.L. Fryer. S.V. Abram is supported by the Department of Veteran Affairs Sierra Pacific Mental Illness Research, Education, and Clinical Center (MIRECC). Dr. Ford is supported by a VA Senior Research Career Scientist award.

## Supplemental Materials

## Abbreviations

*HC*: healthy controls;
*SZ*: schizophrenia subjects;
*ESZ*: early illness schizophrenia subjects;
*CSZ*: chronic illness schizophrenia subjects.

## Additional task description details

The display screen consisted of three reels that populated with fruit symbols with an interstimulus interval of 1 second. There were 12 possible fruit symbols that were equally distributed across conditions (clip art from https://openclipart.org); thus, no individual symbol carried predictive information about reward outcomes. Subjects self-initiated each trial via a button press that triggered the simulated coin drop and slot machine lever pull, after which reels 1-3 (i.e., R1, R2, R3) populated automatically from left to right. After R3 populated, a 10Hz visual checkerboard flickered for 1000 ms followed by text indicating the outcome (i.e., “WIN $1.25” or “LOSE”). Sound effects (e.g., coin dropping audio) and visualizations (e.g., coin insertion, lever pull) enhanced the task’s ecological validity.

The *reward anticipation* phase spanned the population of R1 and R2 and the anticipation of R3. The *reward evaluation phase* began when R3 populated. Total trial time was 6115 ms.

## Additional EEG data denoising procedures

Electro-oculogram (EOG) data were recorded from electrodes placed above and below the left eye and at the outer canthi of both eyes to capture vertical (VEOG) and horizontal (HEOG) eye movements. The Fully Automated Statistical Thresholding for EEG artifact Rejection (FASTER) MATLAB-based toolbox was used to clean the raw EEG epochs (Nolan, Whelan, & Reilly, 2010). Like our previous reports (Hamilton et al., 2019), the FASTER processing approach was modified here between steps 2 and 3 to include canonical correlation analysis (CCA, see below for more detail). The FASTER method employs multiple descriptive measures to search for statistical outliers (± 3 SD from mean). This process included 4 steps: (1) outlier channels were identified and replaced with interpolated values in continuous data, (2) outlier epochs were removed from participants’ single trial set, (3) spatial independent components analysis was applied to remaining trials, outlier components were identified using the ADJUST procedure (Mognon, Jovicich, Bruzzone, & Buiatti, 2011), and data were back-projected without these components, and (4) within an epoch, outlier channels were interpolated. Epochs were time locked to the onset of stimulus and baseline corrected (−100 to 0ms).

Canonical correlation analysis (CCA) was used as a blind source separation technique to remove broadband or electromyographic noise from single trial electroencephalographic (EEG) data, generating de-noised EEG epochs. Our approach is similar to the CCA method described by others (De Clercq, Vergult, Vanrumste, Van Paesschen, & Van Huffel, 2006; Riès, Janssen, Burle, & Alario, 2013), and previously applied to other studies, with some important differences. The method is based on the concepts that true EEG data tend to show high auto-correlation and exhibit power-law scaling (i.e., power is proportional to 1/frequency), but that high frequency random noise in EEG (e.g., muscle artifact, electromyographic (EMG)) tends to show low auto-correlation and violates power-law scaling (i.e., inappropriately high power at higher frequencies relative to low frequencies). The CCA de-noising procedure is performed separately for each subject on the single trial EEG epoch data. For a 3 second epoch, the (S x X) matrix containing the time series of S= 3072 EEG samples (s_1_, s_2_, s_3_ …s_3072_) at each of X=64 scalp electrodes (x_1_, x_2_, x_3_…x_64_) is subjected to a CCA with the (S x Y) matrix containing the s+1 time-lagged series of 3072 EEG samples (s_2_, s_3_, s_4_ …s_3072_, s_3073_, where s_3073_=0) at each of the same 64 electrodes (y_1_, y_2_, y_3_…y_64_). This is the multivariate equivalent of auto-regressive time series correlation. Since both the X and Y vectors each contain 64 electrodes, a total of 64 canonical correlations can be extracted. Each canonical correlation coefficient expresses the correlation of a time series of values representing the weighted sums of the X electrodes with a s+1 time series of values representing the weighted sums of the Y electrodes, with weights chosen to yield the largest canonical correlation that accounts for variance independent of the variance accounted for by all previously extracted canonical correlations. Thus, each canonical correlation coefficient has an associated time series of values that constitutes the canonical variate, X (i.e., each time point has a value that is a linear function of the canonical weights and raw data associated with the 64 electrodes), as well as a similar canonical variate, Y. The current CCA de-noising method only makes use of the set of 64 canonical X variates, one for each of the 64 extracted canonical correlations. When the time series represented by a canonical variate is subjected to a fast Fourier transformation (FFT), the resulting power spectrum can be evaluated to determine whether the canonical variate conforms to the power-law expected from EEG data, in which case it should be retained, or whether it violates the power law as would be expected for high-frequency noise (e.g., EMG contamination), in which case it should be excluded. With this approach, the retained canonical variates are those showing the strongest canonical correlations, whereas the rejected canonical variates are those showing the weakest canonical correlations. The specific criterion used to make these retain/reject decisions is where our CCA denoising approach differs from previously published approaches from other labs (e.g., Riès et al., 2013), but is consistent with our other reports (Hamilton et al., 2019; Kort et al., 2017).

## ERP age-matched HC subgroups

As shown in Table 1, there were significant differences in age between the three groups. As described in the main text, we used an age-adjustment procedure to account for these inherit differences; however, to be certain that our between-group ERP effects were not driven by these demographic differences, we performed follow-up analyses using age-matched HC subgroups. HC subjects were divided into two age-matched using a median split: i) younger HC (*n* = 27) = 23.15 +/− 2.07 years vs. ESZ (*n* = 26) 24.47 +/− 4.01 years, and ii) older HC (*n* = 27) = 44.28 +/− 13.69 years vs. CSZ (*n* = 28) = 44.05 +/− 14.73 years. There were no significant differences in age or gender (all *p* > .10) in either matched group set.

## ERP grand averages

As described previously, there was a significant negative relationship between Age and RewP in HC (Fryer et al., n.d.). We therefore present the grand averages of the RewP difference waveforms in Figure 2, with HC separated into younger and older subgroups (determined via median split as described above). Figure S1 shows the frontal-central scalp distribution of the RewP difference waves, again with the HC separated into younger and older subgroups. We note that the RewP difference score was significantly larger when measured at R3 versus R2 for HC, ESZ, and CSZ (all *p* < .01), consistent with complete reward feedback delivery occurring at R3.

## ERP age-matched subgroup re-analysis

When using age-matched HC subgroups and raw ERP scores, the Group x Condition interactions remained non-significant (younger HC vs. ESZ: F_1,51_ = 0.14, *p* = .71; older HC vs. CSZ: F_1,52_ = 0.04, *p* = .83). However, CSZ had lower SPN amplitudes, overall, relative to older HC; F_1,52_ = 6.15, *p* = .02, indicating reduced anticipation feedback among CSZ when expecting reward or non-reward feedback. (One CSZ subjects was removed who had an SPN value > 3 SD above the mean.)

There was no effect of Group when comparing ESZ and CSZ to age-matched HC subgroups using the unadjusted RewP data (younger HC vs. ESZ: *t*_51_ = 0.73, *p* = .47; older HC vs. CSZ: *t*_53_ = −0.16, *p* = .87).

Finally, we observed the same pattern of LPP group difference, with ESZ showing a heightened near miss response relative to younger HC (F_1,50_ = 8.71, *p* = .005; Figure S2), versus no difference between CSZ and older HC (F_1,52_ = 0.00, *p* = .97). (One ESZ and one CSZ subject were removed who had an LPP values > 3 SD above or below the mean.)

## BrainAGE Model

Results from the HC BrainAGE model are presented in Table S2. We observed a significant negative age-relationship with the RewP, indicating reduced win-related reward responsiveness with age. We also found a positive relationship between age and the LPP average, which could mean that the salience of non-specific motivational factors (wins and losses) increases with age. Further, the SPN difference score’s (possible wins AA – total misses AB) contribution to the model was marginally significant. Taken together, the HC age regression model used to derive estimates of BrainAGE indicated that with increasing age, HCs have reduced early reward evaluation response (RewP), elevated attention to reward outcomes regardless of valence (average LPP) and marginally reduced reward anticipation signaling (SPN).

**Table S1.** Age-adjusted LPP Group x Condition follow-up tests

|  | **t-ratio** | **p-value** | **p-adj** |
| --- | --- | --- | --- |
| **Contrast** |  |  |  |
| *Win Condition (AAA – ABC)* |  |  |  |
| HC wins vs. ESZ wins | 0.77 | .44 | .86 |
| HC wins vs. CSZ wins | 0.41 | .69 | .86 |
| ESZ wins vs. CSZ wins | -.31 | .76 | .86 |
| *Near Miss Condition (AAB – ABC)* |  |  |  |
| HC near misses vs. ESZ near misses | -1.37 | .17 | .47 |
| HC near misses vs. CSZ near misses | 0.34 | .73 | .86 |
| ESZ near misses vs. CSZ near misses | 1.46 | .15 | .47 |
| *Win vs. Near Miss (AAA – ABC vs. AAB – ABC)* |  |  |  |
| ESZ wins vs. ESZ near misses | -3.18 | **.002** | **.02** |
| CSZ wins vs. CSZ near misses | -.09 | .93 | .93 |

Abbreviations: *p-adj*, FDR-adjusted p-values; *HC*, healthy controls; *ESZ*, early illness schizophrenia subjects; *CSZ*, chronic illness schizophrenia subjects.

**Table S2.** HC Brain Age Model

| | $\beta$ (SE) | t-stat | p-value | VIF |
| --- | --- | --- | --- | --- |
| **SPN (AA – AB)** | .24 (.13) | 1.79 | .08 | 1.34 |
| **RewP (AAA – AAA)** | -.53 (.12) | -4.43 | **<.001** | 1.06 |
| **LPP (AAA + AAB)/2** | .41 (.14) | 3.02 | **.004** | 1.38 |
| **LPP (AAA – AAB)** | -.03 (.12) | -0.26 | .79 | 1.03 |

$F_{4,49} = 6.21$ , $p < .001$ , Adjusted $R^2 = .28$
Abbreviations: *VIF*, variance inflation factor; *SPN*, stimulus preceding negativity; *RewP*, reward positivity; *LPP*, late positive potential; *AA*, possible wins; *AB* total misses; *AAA*, wins; *AAB*, near misses.
Mean absolute error = 9.37; Root mean square error = 11.63
Correlation between chronological and predicted ages for HC: $r_{52} = .58$ , $p < .001$

**Figure S1.**
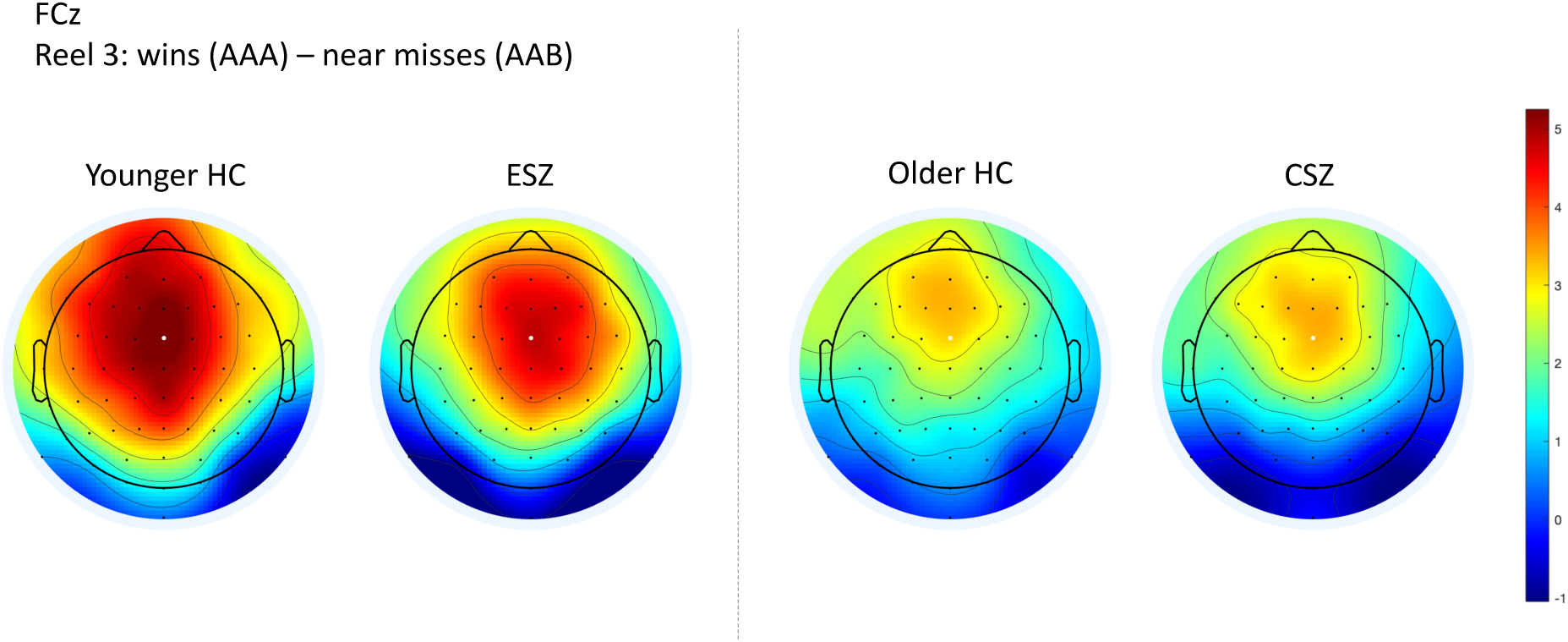
RewP Difference Score Topographical Maps. Topographical maps of condition difference waves at Reel 3 for all groups, wins (AAA) – near misses (AAB).

